# Calcium homeostasis plays important roles in the internalisation and activities of the small synthetic antifungal peptide PAF26

**DOI:** 10.1101/2020.02.14.948786

**Authors:** Akira JT Alexander, Alberto Munoz, Jose F. Marcos, Nick D. Read

## Abstract

Fungal diseases are responsible for the deaths of over 1.5 million people worldwide annually. Antifungal peptides represent a useful source of antifungals with novel mechanisms-of-action, and potentially provide new methods of overcoming resistance. Here we investigate the mode-of-action of the small, rationally designed synthetic antifungal peptide PAF26 using the model fungus *Neurospora crassa*. Here we show that the cell killing activity of PAF26 is dependent on extracellular Ca^2+^ and the presence of fully functioning fungal Ca^2+^ homeostatic/signalling machinery. In a screen of mutants with deletions in Ca^2+^-signalling machinery, we identified three mutants more tolerant to PAF26. The Ca^2+^ ATPase NCA-2 was found to be involved in the initial interaction of PAF26 with the cell envelope. The vacuolar Ca^2+^ channel YVC-1 was shown to be essential for its accumulation and concentration within the vacuolar system. The Ca^2+^ channel CCH-1 was found to be required to prevent the translocation of PAF26 across the plasma membrane. In the wild type, Ca^2+^ removal from the medium resulted in the peptide remaining trapped in small vesicles as in the *Δyvc-1* mutant. It is therefore apparent that cell killing by PAF26 is complex and unusually dependent on extracellular Ca^2+^ and components of the Ca^2+^-regulatory machinery.

**AUTHOR SUMMARY:** Life threatening diseases can be caused when fungi invade human tissues. These invasions often occur when a person’s immune defences are down, often due to treatments for cancer or transplantation. These infections are commonly buried deep within the body and as such are difficult to access and treat. Current medications are often highly toxic to the patient. There is also a worrying rise in drug resistance seen in fungi sampled from patients, with infections effectively untreatable – a death sentence. Antifungal peptides such as PAF26 provide a possible solution by offering a cheap and rapidly produced alternative to conventional drugs. However, unlike antibacterial peptides, little is known about how these small molecules mostly exert their effects and cause death. Using live-cell imaging and deletion mutants, this study provides an analysis of the important roles that Ca^2+^-homeostasis and Ca^2+^-signalling, and possible accompanying vacuolar fusion, play during the dynamic internalization and interaction with and within the fungal cell following PAF26 treatment.

## Introduction

Fungal infections today are among the most difficult diseases to manage in humans(Kohler, Casadevall *et al*. 2014). Fungi collectively kill over 1.5 million people annually which is more than malaria and similar to the death toll from tuberculosis (Brown, Denning *et al*. 2012, Bongomin, Gago *et al*. 2017). Increasing resistance to the limited arsenal of antifungal drugs is a serious concern, especially for Candida and Aspergillus infections, for which the therapeutic options are limited. Overall, there is an urgent need to develop new antifungal strategies to tackle fungal infections (Denning and Bromley 2015, Nicola, Albuquerque *et al*. 2019).

Antifungal peptides (AFPs) and peptide-related molecules are being intensively studied as alternatives for the therapeutic control of pathogenic fungi (Matejuk, Leng *et al*. 2010, Duncan and O’Neil 2013, Rautenbach, Troskie *et al*. 2016, Nicola, Albuquerque *et al*. 2019). A detailed understanding of their antimicrobial mechanisms is of high priority if peptides are to be considered as useful antifungal agents. These studies may also aid the identification of novel targets for antifungal therapy (Muñoz, Gandía *et al*. 2013, Rautenbach, Troskie *et al*. 2016). Furthermore, this mechanistic understanding is guiding the de novo design and modification of natural peptides in order to circumvent their limitations (e.g. instability, toxicity, interactions with other drugs, poor kinetics, resistance mechanisms etc) and thus improve their antimicrobial efficacy (Nicola, Albuquerque *et al*. 2019). Overall, AFPs o□er promising alternatives to standard therapies as anti-infectives and immunomodulatory agents with mechanisms-of-action which are less prone to resistance induction compared to conventional antibiotics (Mahlapuu, Håkansson *et al*. 2016).

PAF26 is a synthetic hexapeptide that has been shown to be highly effective at killing filamentous fungi whilst showing low toxicity against human and bacterial cells (Munoz, Lopez-Garcia *et al*. 2006). Unlike membrane permeabilising antimicrobials, initial investigations into the mode-of-action of PAF26 indicated that it did not directly permeabilise the plasma membrane. Instead, at low fungicidal concentrations, PAF26 has a dynamic antifungal mechanism-of-action that involves at least three stages: peptide interaction with the fungal cell envelope (cell wall and/or plasma membrane), endocytic internalisation and accumulation in the vacuole followed by vacuolar expansion. At a certain point, PAF26 is actively transported out of the vacuole into the cytoplasm, followed by a series of complex and specific intracellular effects whose relationship with the eventual death of the target fungus is still unclear (Muñoz, Marcos *et al*. 2012). Two functional and separate motifs (cationic and hydrophobic domains) in the peptide amino acid sequence have been identified as playing important roles in the antimicrobial mode-of-action (Muñoz, Harries *et al*. 2013). As a result of these studies, PAF26 has been proposed as a simple peptide model for the characterization and study of cationic, cell-penetrating antifungal peptides (Muñoz, Marcos *et al*. 2012, Muñoz, Gandía *et al*. 2013).

From average measurements of fungal cell populations expressing the genetically encoded Ca^2+^-reporter aequorin, cytosolic free Ca^2+^ concentrations ([Ca^2+^]_cyt_) within *Neurospora crassa* spore germlings treated with a low inhibitory concentration of PAF26 (2.5-5 µM) have been shown to exhibit a biphasic increase in response to PAF26, and this biphasic [Ca^2+^]_cyt_ increase is completely dependent on the PAF26 and extracellular Ca^2+^ concentrations. The second phase of the biphasic [Ca^2+^]_cyt_ increase was found to be energy dependent because it was blocked by treatment with the metabolic inhibitor, sodium azide (NaN_3_). This may link PAF26 internalization with actin-dependent endocytosis. Consistent with endocytic internalization playing a key role in PAF26 internalization, is the inhibition by the F-actin inhibitor, Latrucunulin A and a reduced uptake rate in endocytosis by the endocytic mutants *Δrvs-167, Δrvs-161* and *Δrab-5* (Muñoz, Marcos *et al*. 2012, Muñoz, Gandía *et al*. 2013). At high fungicidal concentrations (15 µM), PAF26 killed cells even after NaN_3_ treatment, although in a different manner. Under these conditions, the peptide first bound to the cell envelope as before, but was then observed to directly enter the cytoplasm, indicating passive transport across the plasma membrane at high concentrations (Muñoz, Marcos *et al*. 2012).

In the current research, evidence for Ca^2+^-signalling having a significant role in the PAF26 mode-of-action was analysed by testing and analysing the PAF26 sensitivity of deletion mutants defective in different components of their Ca^2+^-signalling machinery. The pattern and kinetics of peptide interaction, internalization and distribution within the cells of PAF26-resistant mutants were compared with the parental wild type by using fluorescently labelled PAF26 combined with live-cell imaging. The role of external Ca^2+^ in these processes and vacuolar fusion was assessed.

## RESULTS

### Sensitivity of fungal cells to killing by PAF26 is dependent on extracellular Ca^2+^

To confirm that extracellular Ca^2+^ plays a role in determining the sensitivity of *N. crassa* to PAF26 (Muñoz, Marcos *et al*. 2012), and thus Ca^2+^ homeostasis being involved in the mode-of-action of PAF26; the wild type strain was grown in: (1) standard Vögels medium (VM) (0.64 mM Ca^2+^); (2) Ca^2+^ free VM (in which Ca^2+^ had been omitted); and (3) VM with twice the normal concentration of Ca^2+^ (1.28 mM). The IC_50_ values for the wild type under these different media are shown in Fig. 1A. The IC_50_ in standard VM was 3.7 ± 0.6 µM and in VM lacking Ca^2+^ was 6.4 ± 0.9 µM, a significant increase in resistance against PAF26 (*P* ≤ 0.001). VM with 2 X Ca^2+^ had an IC_50_ value of 2.5 ± 0.4 µM, a significant increase in sensitivity to PAF26 (*P* ≤ 0.01) compared with standard VM. These results are consistent with a Ca^2+^-dependent mechanism playing a role in determining the sensitivity of *N. crassa* to PAF26 involving uptake of Ca^2+^ from the external medium.

**Fig. 1.**
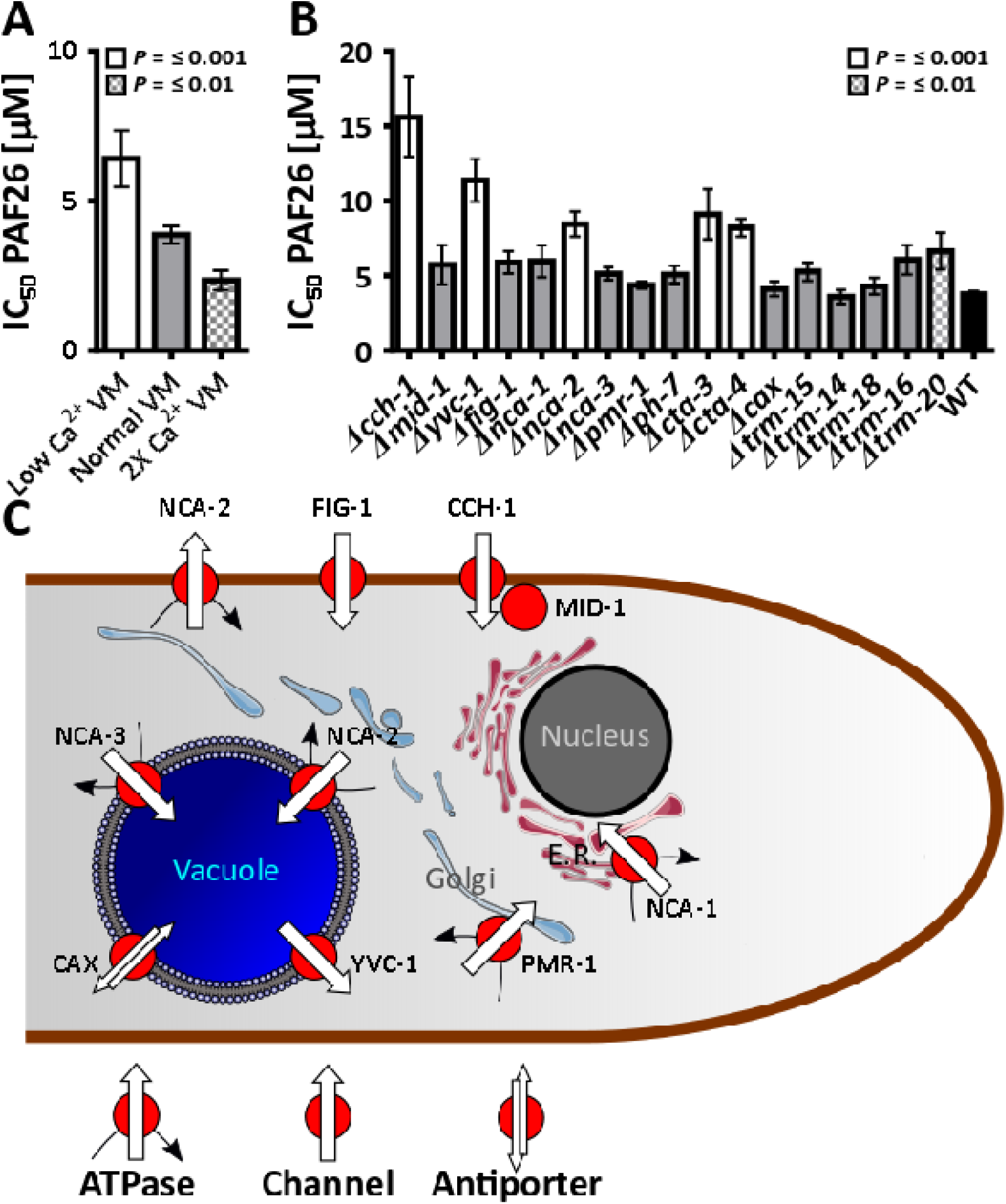
Ca^2+^ plays a significant role in the mode-of-action of PAF26. **(A)** shows the effects of both the removal and increase of free Ca^2+^ in the media on the IC_50_ of PAF26 by fungal cells. Increasing the level lowers the concentration at which PAF26 is effective and removing Ca^2+^ increases the concentration. Both of these results are significant; One way ANOVA with Dunnet’s comparison test: *F*(2,12) = 58.85, *P* =≤ 0.001, R^2^ = 90.75, R^2^(adj) = 89.21. The Low Ca^2+^ VM (M = 6.410 ± 0.941) was significantly different from the Normal VM (M = 3.868 ± 0.288) at *P* = ≤ 0.001 and the 2X Ca^2+^ (M = 2.342 ± 0.331) was significantly different from Normal VM at *P* = ≤ 0.01 **(B)** shows the effects of deleting components of the Ca^2+^ homeostasis machinery on susceptibility to PAF26. 6 were significantly more tolerant than the wildtype; One way ANOVA with Dunnet’s comparison test: *F*(17,106) = 60.13, *P* =≤ 0.001, R^2^ = 90.61, R^2^(adj) = 89.10. *Δcch-1* (M = 15.595 ± 2.649), *Δyvc-1* (M = 11.375 ± 1.419), *Δnca-2* (M = 8.442 ± 0.838), *Δcta-3* (M = 9.099 ± 1.677) and *Δcta-4* (M = 8.212 ± 0.594) were significantly more tolerant than the wildtype (M = 3.756 ± 0.193) at *P* =≤ 0.001, and *Δtrm-20* (M = 6.637 ± 1.234) was significant at *P =* ≤ *0*.*01*. Each assay consisted of 8 technical and 3 biological replicates, figures are representative. The predicted localisations of these is shown in **(C)** based on the localization of their protein orthologs in *S. cerevisiae* yeast cells and vegetative hyphae of *N. crassa* (Wada, Ohsumi *et al*. 1987, Yang and Sachs 1989, Bertl and Slayman 1990, Diamond, Zasloff *et al*. 1991, Antebi and Fink 1992, Iida, Nakamura *et al*. 1994, Lapinskas, Cunningham *et al*. 1995, Levina, Lew *et al*. 1995, Paidhungat and Garrett 1997, Erdman, Lin *et al*. 1998, Kanzaki 1999, Benito, Garciadeblas *et al*. 2000, Locke, Bonilla *et al*. 2000, Muller, Locke *et al*. 2001, Palmer, Zhou *et al*. 2001, Courchesne 2002, Courchesne and Ozturk 2002, Denis and Cyert 2002, Gupta, Ton *et al*. 2003, Kaiserer, Oberparleiter *et al*. 2003, Muller, Mackin *et al*. 2003, Zhou, Batiza *et al*. 2003, Zelter, Bencina *et al*. 2004, Brand, Shanks *et al*. 2007, Hallen and Trail 2008, Lew, Abbas *et al*. 2008, Benito, Garciadeblás *et al*. 2009, Bormann and Tudzynski 2009, Bowman, Draskovic *et al*. 2009, Binder, Chu *et al*. 2010, Bowman, Abreu *et al*. 2011, Cavinder, Hamam *et al*. 2011).

### PAF26 requires a functional [Ca^2+^]_cyt_ homeostatic machinery in order to kill fungal cells

To assess whether Ca^2+^-homeostasis and signalling play a role in the mode-of-action of PAF26; 24 homokaryotic deletion mutants of genes encoding proteins predicted to be components of the *N. crassa* Ca^2+^-homeostatic machinery were screened for increased PAF26 tolerance / sensitivity. The predicted locations of these proteins in the fungal cell are shown in Fig. 1C; The results of the screen are in Table 1.

**Table 1:**
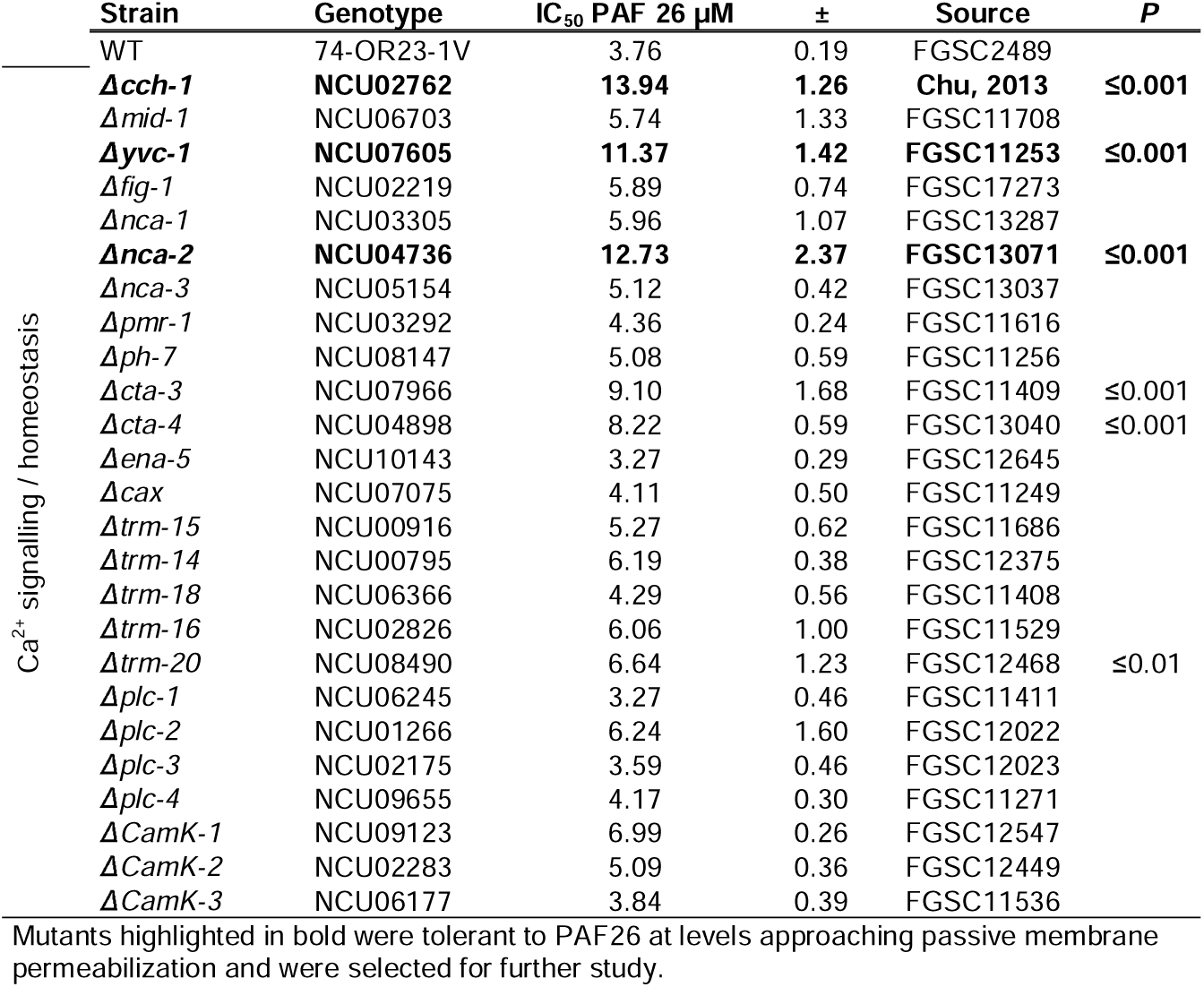
The half maximal inhibitory concentration (IC_50_) of PAF26 in Ca^2+^ signalling/homeostasis mutants.

The first group of mutants screened were defective in the following Ca^2+^-channel proteins: CCH-1 and MID-1, which are components of the high affinity Ca^2+^-influx system (HACS); FIG-1, which forms the low-affinity Ca^2+^-influx system (LACS); and, the vacuolar Ca^2+^ channel, YVC-1. Two of these mutants (Δ*cch-1* and Δ*yvc-1*) exhibited significantly increased tolerance against PAF26 (*P* ≤ 0.001) when compared with the wild type (Fig. 1B). The IC_50_ values of Δ*cch-1* and Δ*yvc-1* were both above the level (∼ 10 µM) at which PAF26 directly permeabilizes the plasma membrane of *N. crassa* wild type cells (Muñoz, Marcos *et al*. 2012).

*N. crassa* has eight Ca^2+^ ATPases (Zelter *et al*., 2004), for which seven deletion mutants were available as homokaryons for screening: Δ*nca-1*, Δ*nca-2*, Δ*nca-3*, Δ*pmr-1*, Δ*ph-7*, Δ*cta-3* and Δ*cta-4. Δnca-2*, Δ*cta-3* and Δ*cta-4* exhibited significantly increased tolerance against PAF26 (*P* ≤ 0.001) when compared with the wild type (Fig. 1B); with *Δnca-2* approaching inhibitory levels of PAF26 that directly permeabilise the membrane in the wild type.

Homokaryotic deletion mutants of six of the eight predicted *N. crassa* antiporters involved in Ca^2+^ transport (Zelter *et al*., 2004) were available for screening: Δ*cax*, Δ*trm-14*, Δ*trm-15*, Δ*trm-16*, Δ*trm-18*, and Δ*trm-20* (Fig. 1B, Table 1). Δ*trm-20* showed significantly greater tolerance (*P* ≤ 0.01) to PAF26 compared with the wild type although this was less than a two-fold increase (Fig. 1B).

### PAF26 is taken up by wild type macroconidia into vacuoles which fuse and then lyse before releasing the peptide into the cytoplasm

In order to compare the dynamic pattern of peptide-cell interactions between the wild type and the mutants, PAF26 labelled with the fluorophore TAMRA (TMR-PAF26) was imaged. Fluorescent labelling has been previously employed to monitor, using confocal live-cell imaging, the dynamic localization of PAF26 or related peptides in a range of fungi. We are confident that the dynamic pattern of TMR-PAF26 staining provides a faithful localization of the unlabelled peptide in fungal cells and labelled/unlabelled peptides have similar IC_50_ values (Muñoz, Marcos *et al*. 2012, Muñoz, Gandía *et al*. 2013, Muñoz, Harries *et al*. 2013). Previously it has been shown that wild type macroconidia treated with a low fungicidal concentration of FITC-PAF26 (2.5 µM) exhibit a similar time-dependent staining pattern as germlings treated with the same concentration of PAF26 (Muñoz, Marcos *et al*. 2012). A similar localization pattern of TMR-PAF26 interaction, internalization and distribution was observed with ungerminated macroconidia to that of FITC-PAF26. The use of macroconidia allowed a basic monitoring of cell volume, surface area and developmental state in relation to possible variations in morphogenesis and staining patterns.

### The dynamic pattern of TMR-PAF26 in Ca^2+^ homeostasis mutant staining provides insights into the localization and roles of the defective mutant proteins

In order to gain more insight into which stages of the PAF26 interaction, uptake and distribution process the CCH-1, YVC-1 and NCA-2 proteins influence, the time-dependent staining by TMR-PAF26 in each of the *Δcch-1*, Δ*yvc-1*, and *Δnca-2* mutants were compared with that in the wild type. As a baseline for the comparison, four staining patterns were identified in the wild type macroconidia that could be readily quantified at 30 min intervals following treatment with 3.5 µM TMR-PAF26. These staining patterns related to different events in the time-dependent uptake and distribution to different organelles which precedes eventual cell death. They were: (a) Accumulation in multiple small vesicles. These were predicted to be mostly endosomes typically up to ∼ 1 µm in width. (b) Accumulation in 1-3 large vacuole(s) typically greater than ∼ 1.5 µm in width. (c) Accumulation in the cytoplasm and excluded from vacuoles. This is the stage just after the peptide has been released out from vacuoles. (d) Accumulation in both the cytoplasm and vacuoles. This is the stage after which the intracellular membranes have become permeabilized and thus represents commitment to cell death, as previously reported (Muñoz, Marcos *et al*. 2012).

In the wild type macroconidia (Figs. 2B and 3A), this quantitative analysis showed that after treatment for 30 min with TMR-PAF26, all the macroconidia had interacted with TMR-PAF26 and many were beginning to internalise the peptide into small vesicles. After 60 min, many of the conidia had already undergone vacuolar expansion and some had exported the peptide into the cytoplasm, and this correlated with a transient reduction in the percentage of cells with peptide only localized within small vesicles. After 60 min, 40% of the cells had actively transported the peptide out of their vacuoles and into the surrounding cytoplasm. In another 40% of the cells at the 60 min time point, vacuolar membrane permeabilisation had occurred and the peptide had equilibrated throughout the cells. By 120 min, 57% of the cells were permeabilised (Figs. 2B and 3A).

**Fig. 2.**
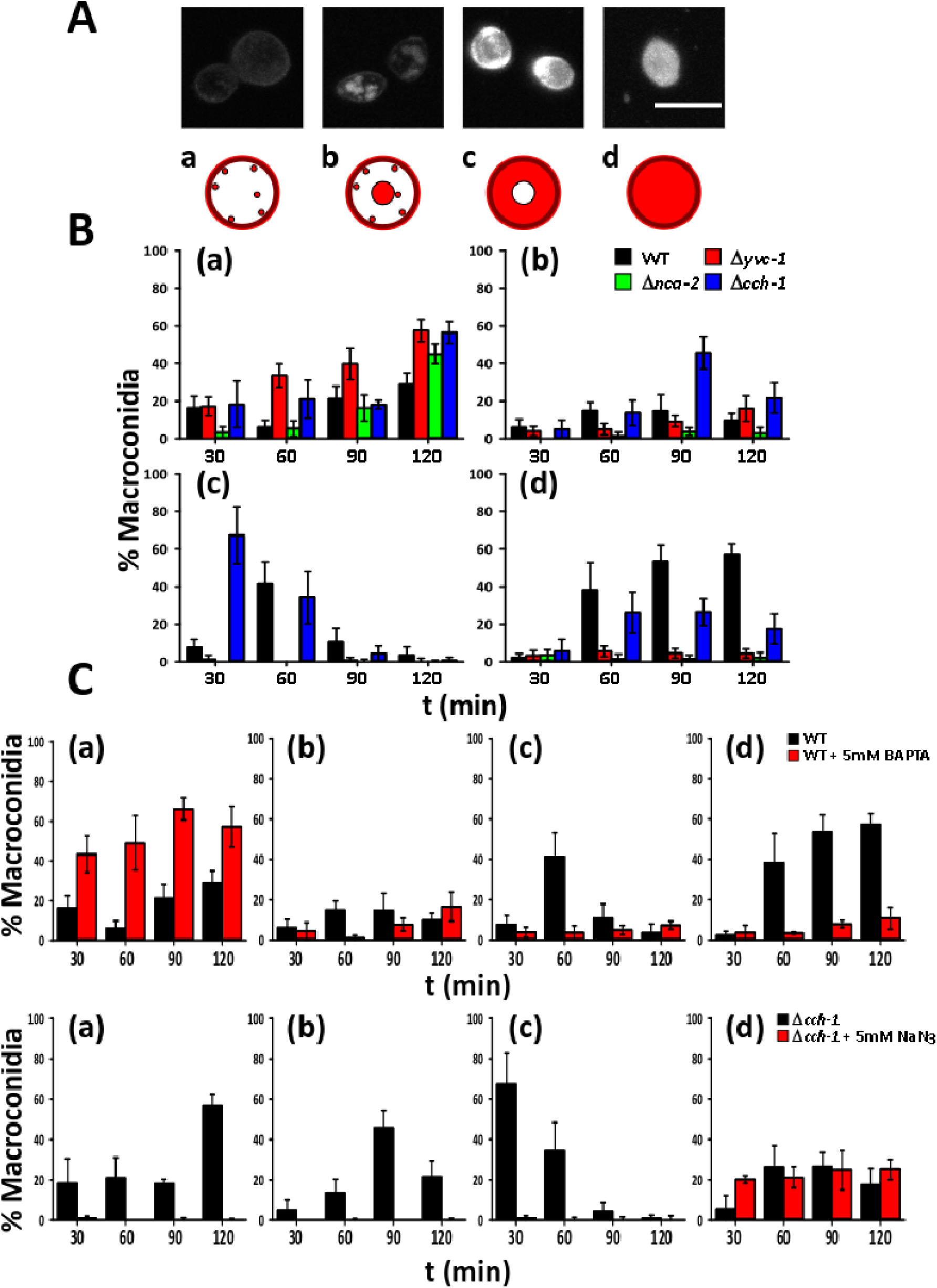
Graphical representation showing that the interaction between PAF26 and macroconidia is altered through deletion of Ca^2+^-channels and pumps. (**A)** shows maximum intensity projections from a Z series captured using confocal microscopy. Initially, the peptide appears at the cell envelope and within small vesicles (a). The peptide is then accumulated in larger vesicles and vacuoles (b) until there appears a lower number of larger vacuoles (c) which will eventually fuse into a single vacuole which then releases the peptide into the cytoplasm (d). Scale bar is 5 µm. **(B)** shows quantification of this interaction over time for the deletion mutants compared to the wild type; there are marked changes in the localisation of the peptide during the time course. **(C)** shows the interaction between PAF26 and macroconidia is Ca^2+^-dependent and energy dependent. This is demonstrated by the significant delay in the uptake of PAF26 in the wild type when extracellular free Ca^2+^ is removed using BAPTA, at top. The peptide remains trapped in small vesicles and there is little transport into the vacuolar system and consequently a reduction in internal membrane permeabilisation. The lower figure shows the effect of the removal of cellular free energy on the *Δcch-1* mutant using the metabolic inhibitor NaN_3_. There is no uptake of PAF26 into the cell and therefore the direct translocation across the plasma membrane must be an energy dependent process. N = 100 macroconidia minimum per field of view, 10 fields of view per timepoint, repeated at least twice.

**Fig. 3.**
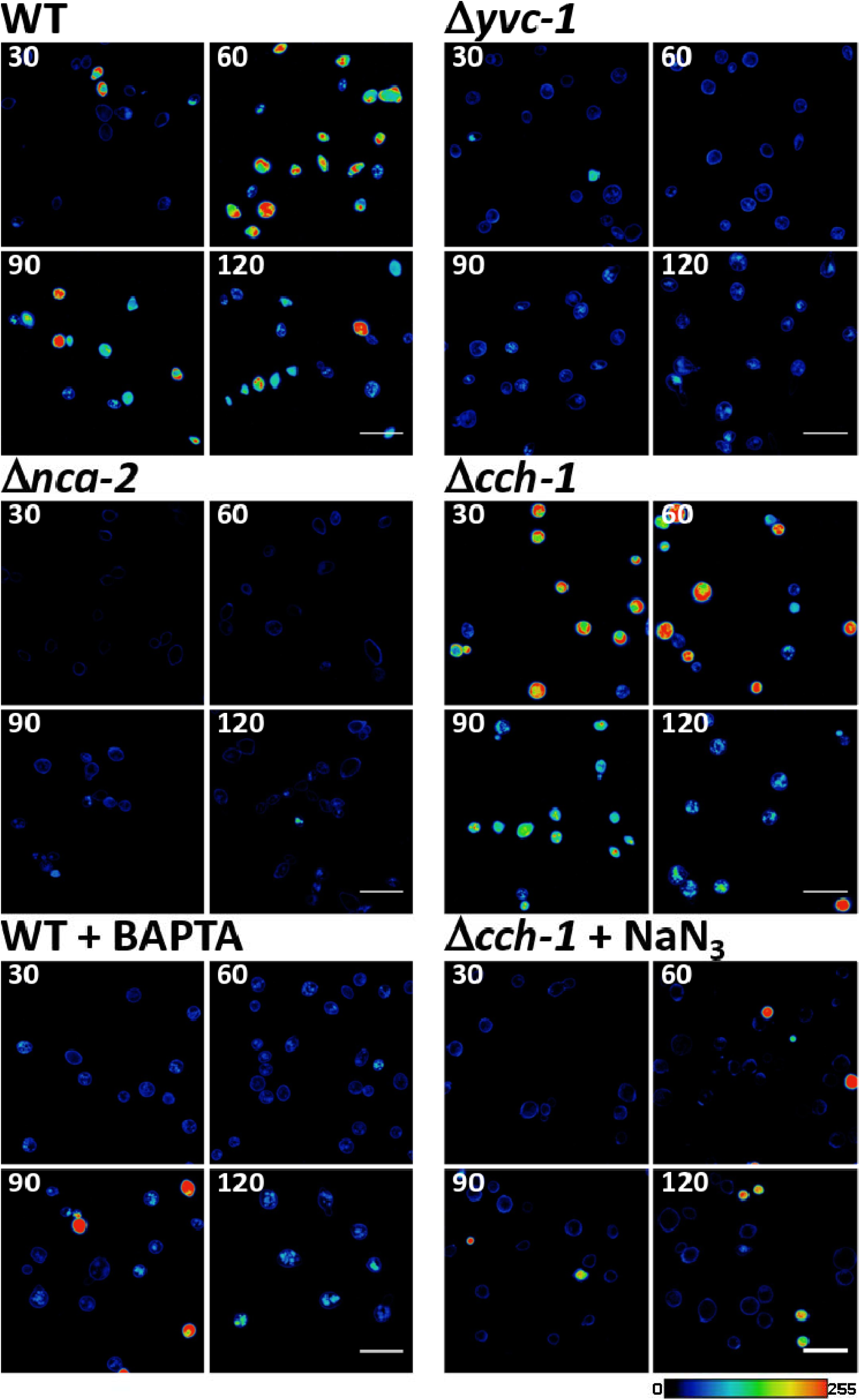
The interaction with PAF26 is altered through deletion of Ca^2+^-channels and -pumps as shown by visualisation of wild type and mutant conidia stained with 3.5 µM TAMRA labelled PAF26. These figures show maximum intensity projections of Z stacks false coloured with the ‘physics’ lookup table to highlight the different fluorescent intensities measured. (**A)** The wild type conidia undergo a sequence of events which ultimately results in cell death (Muñoz, Marcos *et al*., 2012). (**B**) The *Δyvc-1* mutant retains the peptide in small vesicles, without the formation of a large central vacuole, as in the wild type. (**C**) The *Δnca-2* mutant appears much reduced in peptide interaction, with very few conidia showing the peptide bound to the envelope. (**D**) The process is reversed in the *Δcch-1* mutant, where the peptide first appears within the cytoplasm and is pumped into small vesicles which then appear to fragment. No conidia were seen to completely lose loaded peptide. The internalisation process is Ca^2+^ dependent and energy-dependent, as chelation of Ca^2+^ with BAPTA (**E**), and treatment with Sodium azide (NaN_3_) (**F**), significantly reduced uptake of TAMRA-PAF26. 1000 individual macroconidia per timepoint per strain. Assay was repeated twice and images are representative. Scale bar is 20 µm.

### YVC-1 is required for transport of PAF26 from vesicles to vacuoles

When the Δ*yvc-1* mutant was treated with the labelled peptide (Fig. 2B, 3B), it rapidly appeared at the cell envelope. Unlike with the wild type, the peptide was internalised into small vesicles at a more-or-less linear rate with much remaining trapped at the cell surface. Accumulation of the peptide in vacuoles was much slower and reached a maximum of 6% of the cells with this localization pattern after 120 min. Virtually no cells were observed with peptide exclusively in the cytoplasm at the three time points of measurement over the 120 min by which time only 5% of cells were internally permeabilised. After 120 min, 79% of the Δ*yvc-1* cells had taken up TMR-PAF26 compared with 100% in the wild type. Thus, less TMR-PAF26 was taken up by Δ*yvc-1* cells, most accumulated in small vesicles and few cells had internal membrane permeabilisation after 120 min (5% compared with 57% in the wild type). These results are consistent with YVC-1 being required for the transport of PAF26 from vesicles to vacuoles.

### NCA-2 is required for the interaction of PAF26 and the cell envelope

When the Δ*nca-2* mutant was treated with TMR-PAF26 (Fig. 2B and 3C) the cell envelopes of the macroconidia were visibly less fluorescent than that of the wild type. Thus, the affinity of TMR-PAF26 for the cell envelope of this mutant seemed to be reduced and its staining was delayed compared with that of the wild type. The Δ*nca-2* macroconidia also internalized TMR-PAF26 at a far reduced rate compared with the other mutants and the peptide mostly became trapped within the small vesicles, resulting in virtually no TMR-PAF26 being taken up by vacuoles and a correspondingly extremely small percentage (∼ 2%) of the cells having permeabilised internal membranes after 120 min (Fig. 2B).

### CCH-1 plays a key role in the energy- and Ca^2+^-dependent internalisation of PAF26 by fungal cells

The pattern of TMR-PAF26 internalization and intracellular transport within macroconidia of the *Δcch-1* mutant appeared to be opposite to that in the wild type (Fig. 3D). After treatment for 30 min the peptide was primarily localized in the cytoplasm (in 67% of cells) and to a much lesser extent in small vesicles (∼ 18% of cells) and to a very low level in large vacuoles (∼ 5% of cells). Between 30 and 90 min, TMR-PAF26 was removed from the cytoplasm and increased in amount in the small vesicles and large vacuoles. Between 90 and 120 min the large vacuoles containing TMR-PAF26 are fragmenting into smaller vesicles. This was reflected by a dramatic increase in the number of stained small vesicles (Fig. 2B).

The overall rate of internalisation of TMR-PAF26 by *Δcch-1* macroconidia was faster than in the wild type because in the former it was initially taken up directly into the cytoplasm whilst in the latter it first appeared intracellularly in small vesicles that are presumed to be mostly endosomes. It seems unlikely that TMR-PAF26 is taken up by means of non-specific permeabilization of the plasma membrane because this would likely have resulted in rapid cell death. Furthermore, 30 min after the addition of TMR-PAF26 it was clear that the vacuolar membrane had not become permeabilized because the peptide was excluded from the vacuoles of cells in which the cytoplasm was fluorescent.

In order to clarify whether the passage of PAF26 across the plasma membrane into the cytoplasm of the *Δcch-1* mutant was a result of passive translocation or active uptake, macroconidia were pre-treated with the metabolic inhibitor NaN_3_ (Muñoz, Marcos *et al*. 2012) at a concentration of 5 µM for 15 min before the addition of TMR-PAF26 and subsequent imaging and quantification of localization over 120 min (Figs. 2C and 3F).

In the presence of NaN_3_, over the whole 120 min period of incubation with TMR-PAF26, the peptide remained bound to the cell envelope and was not internalized by most of the macroconidia. Clearly the metabolic inhibitor NaN_3_ had almost completely abolished the movement of TMR-PAF26 across the plasma membrane indicating that the uptake of the peptide into the cells of the *Δcch-1* mutant is an ATP-dependent process. The results also showed that the rate of internal membrane permeabilisation, whilst faster over the first 30 min, remained similar to that of the non-azide treated *Δcch-1* cells; after 120 min ∼ 25% of the cells were fluorescent throughout the cell. These results are consistent with the uptake of TMR-PAF26 by this subpopulation of *Δcch-1* macroconidia being energy independent as a result of a passive process.

These results are consistent with the normal endocytic internalization of PAF26 being dependent on the Ca^2+^ channel protein, CCH-1. As CCH-1 appears to initiate PAF26 internalization by endocytosis (see previous section), this suggests that the uptake of Ca^2+^ from the external medium may be mediated by this Ca^2+^ channel, which has previously been reported to be immunolocalized to the plasma membrane (Locke *et al*., 2000). Furthermore, we had previously shown that removal of Ca^2+^ from VM made macroconidia more resistant to being killed by PAF26. To attempt to mimic the effect of *cch-1* deletion in the wild type, Ca^2+^ was removed from the external medium by the addition of the Ca^2+^ chelator BAPTA (5 mM) 30 min prior to the TMR-PAF26 treatment (Figs. 2C and 3E). Rather than preventing the entry of the peptide, the BAPTA treatment had the unexpected effect of trapping the peptide in small vesicles. Over the 120 min period very little TMR-PAF26 had localized in large vacuoles, in the cytoplasm and only ∼ 11% of the macroconidia had fluorescence throughout the cell. The phenotype of the *Δcch-1* mutant is consistent with the CCH-1 protein playing a key role in the energy- and Ca^2+^-dependent internalisation of PAF26 by fungal cells.

### The [Ca^2+^]_cyt_ response during PAF26 treatment is disrupted in the [Ca^2+^]_ext_ tolerant mutants

The wild type had previously been shown to undergo a dose dependent biphasic rise in [Ca^2+^]_cyt_ upon addition of PAF26 at concentrations between 0.8 and 2.0 µM (Muñoz, Marcos *et al*. 2012). The experiment was repeated here but with over a concentration range of 1.25-10 µM PAF26. For all measurements, the unstimulated [Ca^2+^]_cyt_ resting level was measured for 50 sec, in the wild type this was calculated to be 0.05 ± 0.02 µM (Fig. 4A), before the peptide was added at time 0. After treatment with a low final concentration of PAF26 (1.25 µM), an immediate increase in [Ca^2+^]_cyt_ to 0.19 ± 0.01 µM was observed and this was sustained throughout the measurement period (1,200 sec) and slightly increased towards the end of this period (Fig. 4B). When the PAF26 added was at a final concentration of 2.5 µM, which is close to its IC_50_ value, the initial increase in [Ca^2+^]_cyt_ was slightly greater (to 0.22 ± 0.03 µM) (Fig. 4B). Again, this was followed by a period of sustained [Ca^2+^]_cyt_ increase, but there was also a more pronounced exponential increase in [Ca^2+^]_cyt_ which began at ∼ 900 sec after treatment. With the higher dose, there was also an increase in the standard deviations of the measurements with time. This is due to aequorin consumption during the course of the experiment, resulting in less sensitivity after prolonged exposure to high [Ca^2+^] (note that luminescence measurements from 6 wells are averaged per time point). When the wild type was treated with 5 µM PAF26, which was above its IC_50_ value, the [Ca^2+^]_cyt_ response followed the same general pattern but with a much larger initial [Ca^2+^]_cyt_ increase to 0.32 ± 0.02 µM and a shorter period of sustained increase before the [Ca^2+^]_cyt_ increase became exponential at ∼ 430 sec (Fig. 4B). Furthermore, after 590 sec following peptide treatment, [Ca^2+^]_cyt_ spiking was observed. The [Ca^2+^]_cyt_ response was further accentuated after treatment with a very high dose (10 µM) of PAF26 resulting in a rapid initial rise in [Ca^2+^]_cyt_ to 0.54 ± 0.03 µM, followed by an immediate exponential increase in [Ca^2+^]_cyt_. This in turn was followed by the [Ca^2+^]_cyt_ spiking and then a sudden drop in [Ca^2+^]_cyt_ after ∼ 220 sec, which then exponentially increased and subsequently underwent spiking again (Fig. 4B).

**Fig. 4.**
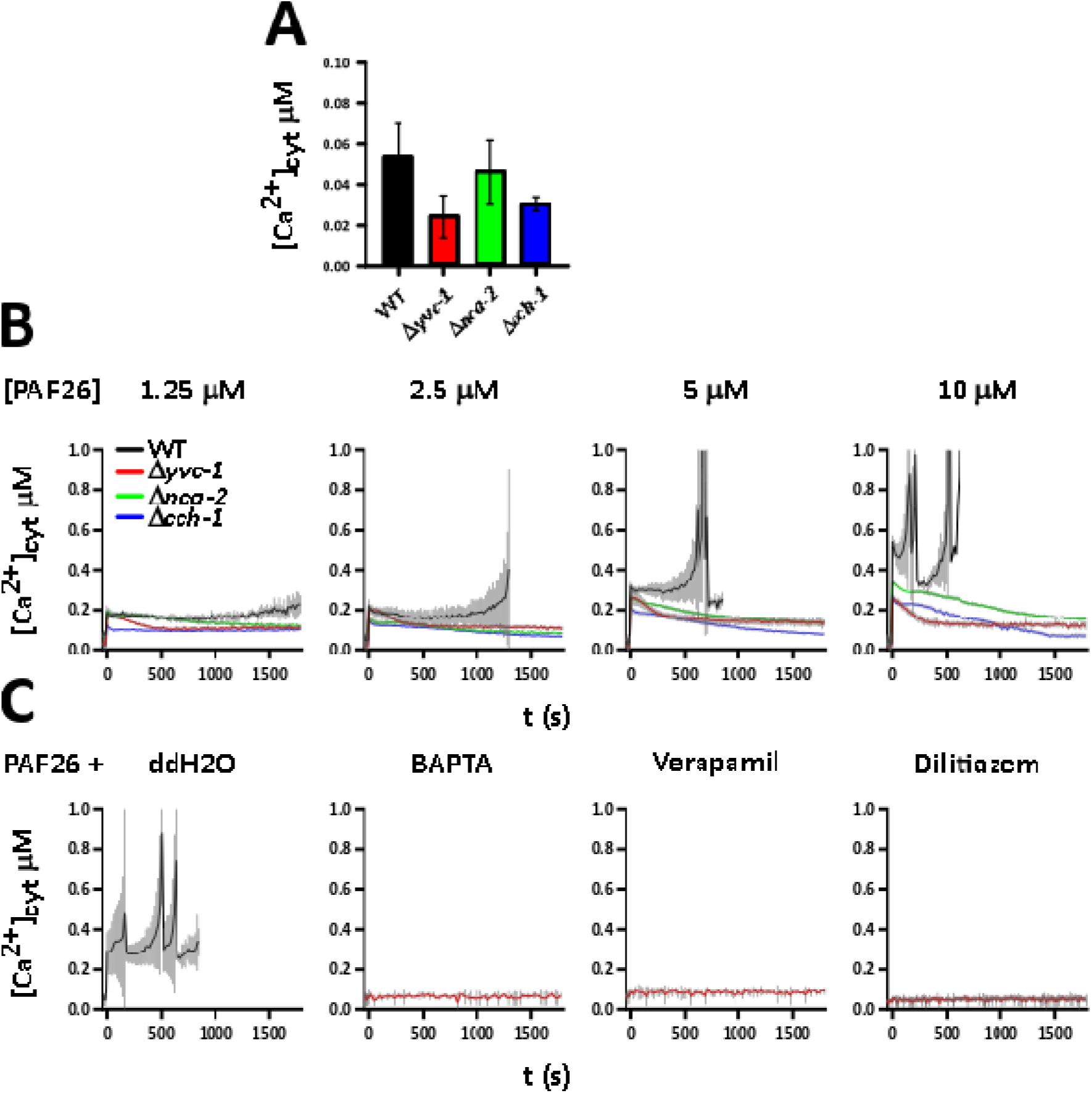
The [Ca^2+^]_cyt_ response to PAF26 is altered by deletion of Ca^2+^ channels and pumps. (**A**) shows the unstimulated resting level of the wildtype and the Ca^2+^ homeostatic mutants: the resting level in *Δyvc-1* and *Δcch-1* were significantly different from the wild type. One way ANOVA with Dunnet’s comparison test: *F*(3,108) = 32.83, *P* =≤ 0.001, R^2^ = 47.70, R^2^(adj) = 46.24. (**B**) shows the [Ca^2+^]_cyt_ response to PAF26 by population wide measurements using the genetically encoded reporter aequorin supplied with an excess of coelenterazine. The [PAF26] µM is shown above each measurement series and was added at time 0, baseline measurements were recorded for 50s before the addition of the peptide. (**C**) shows that this response in the wild type is both dependent on external Ca^2+^ and on the presence of an external Ca^2+^ channel, the biphasic response to 5µM PAF26 (seen in the ddH_2_O control) is completely abolished following chelation of Ca^2+^ with BAPTA or treatment with L-Type Ca^2+^ channel blockers. All measurements are averages of 6 individual wells in a 96 well plate, with [Ca^2+^]_cyt_ calculated following quenching of the remaining aequorin using EtOH and Ca^2+^. Error bars are standard deviations. All experiments were repeated a minimum of three times and the above figures are representative.

The unstimulated resting level of [Ca^2+^]_cyt_ in the Δ*yvc-1* mutant was measured as 0.02 ± 0.01 µM (Fig 4A), significantly different at *P* >0.01. The [Ca^2+^]_cyt_ response of the Δ*yvc-1* strain to PAF26 was markedly different to that of the wild type. Whilst there was a similar initial immediate, dose dependent increase in [Ca^2+^]_cyt_, there was no second exponential increase in [Ca^2+^]_cyt_, which was particularly evident after treating the wild type with PAF26 at 2.5 µM or above (Fig. 4B). Instead the Δ*yvc-1* mutant underwent a slight reduction in [Ca^2+^]_cyt_ followed by a sustained, more-or-less constant [Ca^2+^]_cyt_ level that was significantly raised compared with the unstimulated resting levels recorded prior to treatment. Thus the Δ*yvc-1* mutant lacked the second [Ca^2+^]_cyt_ increase of the biphasic [Ca^2+^]_cyt_ response of the wild type, which is consistent with this second [Ca^2+^]_cyt_ increase resulting from the release of Ca^2+^ from the vacuole. The resting level of [Ca^2+^]_cyt_ in the Δ*nca-2* mutant was measured at 0.05 µM ± 0.02, Fig 4A, and was not significantly different from the wild type. In general terms, the Δ*nca-2* mutant showed a similar response to PAF26 as did the Δ*yvc-1* mutant. However, the Ca^2+^ signatures of the Δ*nca-2* and Δ*yvc-1* mutants were clearly but not dramatically different from each other; there was a more pronounced dose dependent [Ca^2+^]_cyt_ increase in the Δnca-2 strain compared with that in Δ*yvc-1* in response to PAF26 (Fig. 4B). The resting level of [Ca^2+^]_cyt_ in the Δ*cch-1* mutant was 0.03 ± 0.00 µM Fig. 4A, significantly different to the wild type at *P* > 0.01. Interestingly, deletion of *cch-1* does not inhibit the initial increase in [Ca^2+^]_cyt_ after PAF26 addition, surprising given the lack of evidence for another Ca^2+^ channel at the cell surface. The Ca^2+^ signatures of the Δ*cch-1* mutant in response to different concentrations of PAF26 were broadly similar to those of the Δ*yvc-1* and Δ*nca-2* mutants (Figs. 4B). Thus the Δ*cch-1* mutant also showed an initial [Ca^2+^]_cyt_ increase but lacked the second [Ca^2+^]_cyt_ increase of the typical biphasic response of the wild type to PAF26. These results suggest there may be an unidentified L-type channel within the plasma membrane, not CCH-1, being responsible for the initial [Ca^2+^]_cyt_ increase in response to PAF26.

The influence on the [Ca^2+^]_cyt_ response to 5.0 µM PAF26 of the wild type following pre-treatment with 5 mM BAPTA (to chelate extracellular Ca^2+^) or with the L-type Ca^2+^ channel blockers diltiazem or verapamil (both at 5 mM), was analysed (Fig. 4C). A 15 min pre-treatment with ddH_2_O (the control) caused a biphasic increase in [Ca^2+^]_cyt_ whilst all of the other treatments prevented any significant [Ca^2+^]_cyt_ increase (Fig. 2). These results are consistent with the biphasic [Ca^2+^]_cyt_ increase being dependent on extracellular Ca^2+^ as reported before (Muñoz, Marcos *et al*. 2012) and on L-type Ca^2+^ channel activity. The only known L-type Ca^2+^ channel in *N. crassa* is CCH-1(Zelter, Bencina *et al*. 2004). These results also suggest that the second component of the biphasic [Ca^2+^]_cyt_ increase is dependent on the first [Ca^2+^]_cyt_ increase.

### Imaging using the fluorescent reporter GCaMP6 shows PAF26 does not cause a dose dependent rise in [Ca^2+^]_cyt_

In order to investigate whether PAF26 was indeed causing a dose dependent rise in [Ca^2+^]_cyt_, conidia expressing GCaMP6s were imaged during PAF26 treatment using widefield fluorescence microscopy. Individual macroconidia were then isolated from the background in FIJI and fluorescence measured over time. As Figure 5A shows, the individual responses are far more varied than the measurements using aequorin suggested. In the middle of the timecourse, the moment the PAF26 reaches the field of view and interacts with the cells can be clearly seen in a sudden increase in Ca^2+^ spiking in all the individuals. Given that the aequorin measurements are across a whole population, it is not hard to see how the cumulative effects of an increase in Ca^2+^ spikes and waves could be interpreted as a dose dependent rise in [Ca^2+^]_cyt_.In order to determine whether there was a predictable response to PAF26 addition, several repeated experiments were run in which the peptide was added and fluorescence intensity recorded. No distinct pattern or regularity was found within any of the data sets, except for an increase in Ca^2+^ signalling after PAF26 addition. As the only noticeable response, the amplitude and frequency of these Ca^2+^ spikes was quantified. Conidia were treated either with ddH_2_O or ddH_2_O containing 3.5 µM PAF26. A second set were pre-treated with 5 mM BAPTA before ddH_2_O or PAF26 treatment. The conidia treated with ddH_2_O have a relatively low frequency of Ca^2+^ spiking with the mean being 4 spikes over the course of the measurement. This is consistent with our findings as to the rate of Ca^2+^ spiking in germinating and fusing conidia (Read, unpublished). When the cells are treated with PAF26 however, there is a marked increase in both frequency and amplitude. The mean frequency of the Ca^2+^ spikes increases to 19 and the amplitude increases to around 6 times the resting level from 4 (Figure 5B). Both of these results are significant at *P* < 0.01. When the conidia are pre-treated with BAPTA however, all Ca^2+^ spiking ceases completely. Whilst there were no apparent recurring patterns in the spiking events, observations appeared to show that they correspond to vesicles and vacuoles coming into close proximity. A Ca^2+^ trace is shown in Fig. 5C and images are shown in Fig. 5D. The images shown in Fig. 5D are taken every five seconds from the timeframes indicated by the black arrows in Fig. 5C. The first Ca^2+^ spike marked by *, appears to correspond with the meeting of the two vacuoles at the lower part of the image, whether they fuse or not is unclear. In the images corresponding to the second spike marked with *, the fusion of two vacuoles is clear at the top of the image.

**Figure 5.**
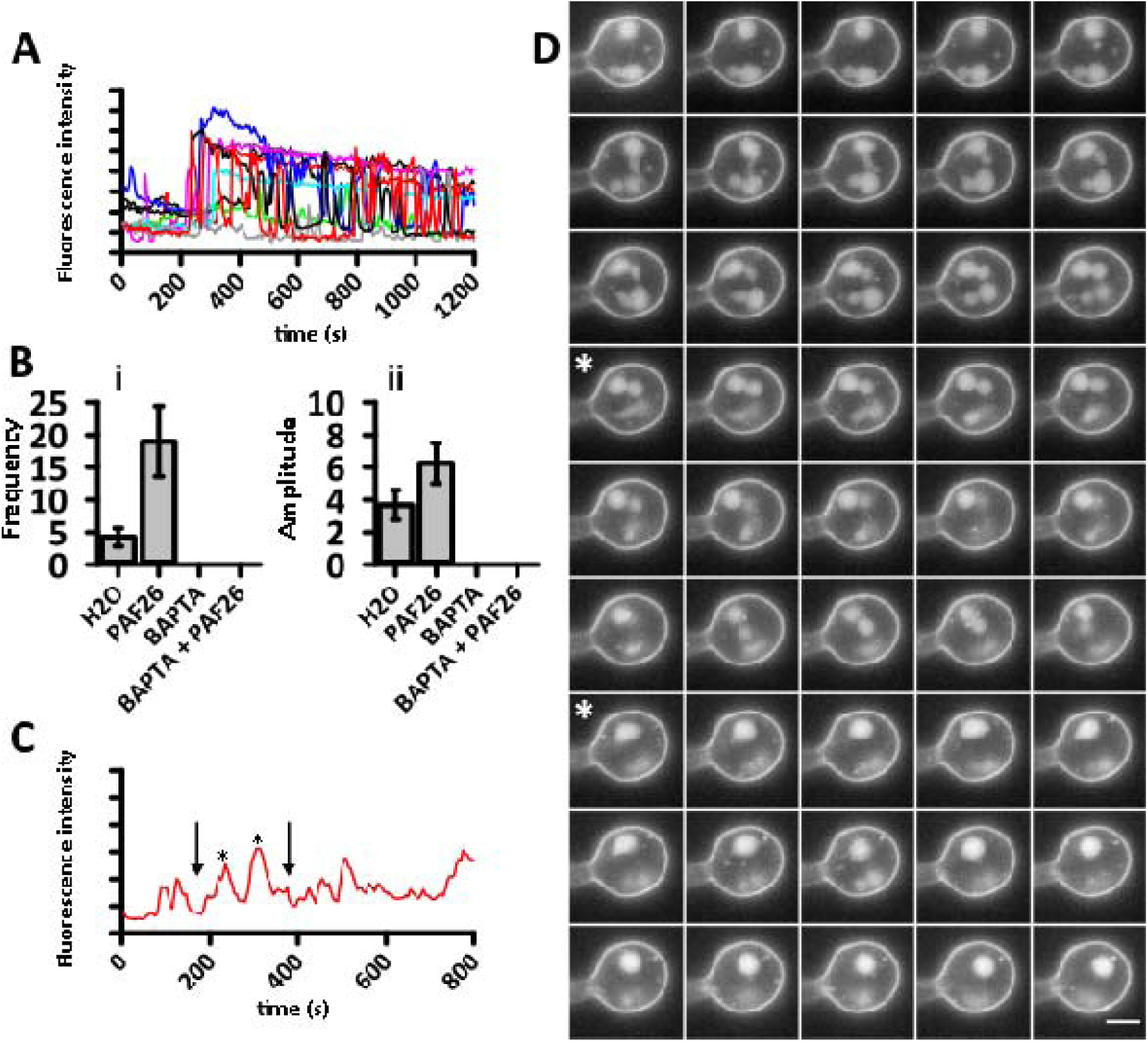
PAF26 does not cause a dose dependent rise in [Ca^2+^]_cyt_. (**A**) shows the relative fluorescence intensity of several macroconidia expressing GCaMP6s during PAF26 treatment. Fluorescence values are normalised and given as arbitrary values. After the addition of PAF26 there is a marked increase in the amplitude and frequency of Ca^2+^ spiking. (**B**) Quantification of this revealed an actual increase in spiking from an average of 4 spikes per 30 min to 19 spikes per 30 min (i). The spiking was entirely dependent on the presence of external Ca^2+^ as chelation with BAPTA reduced the frequency of spiking to 0. The amplitude of the individual spikes also increased from 4 to 6 arbitrary units (ii). (**C**) these spikes often appeared to correspond to the meeting of vesicles and vacuoles within the cell; the images captured between the two arrows are shown in (**D)** which shows widefield fluorescence images captured every 5 s. The two spikes marked by * in C appear to correspond to vacuolar fusion events. Scale bar = 2µm.

### YVC-1 is localized in vacuoles and CCH-1 is both plasma membrane and vacuolar localized

In *N. crassa*, CCH-1 and YVC-1 have not yet been definitively localised while NCA-2 has previously been localised to the vacuole and plasma membrane in mature hyphae (Bowman, Draskovic *et al*. 2009). We conducted experiments to localize NCA-2 tagged with RFP in macroconidia of *N. crassa* and the data showed labelling of the vacuoles, but no obvious signal at the plasma membrane (Fig. 6).).

**Fig. 6.**
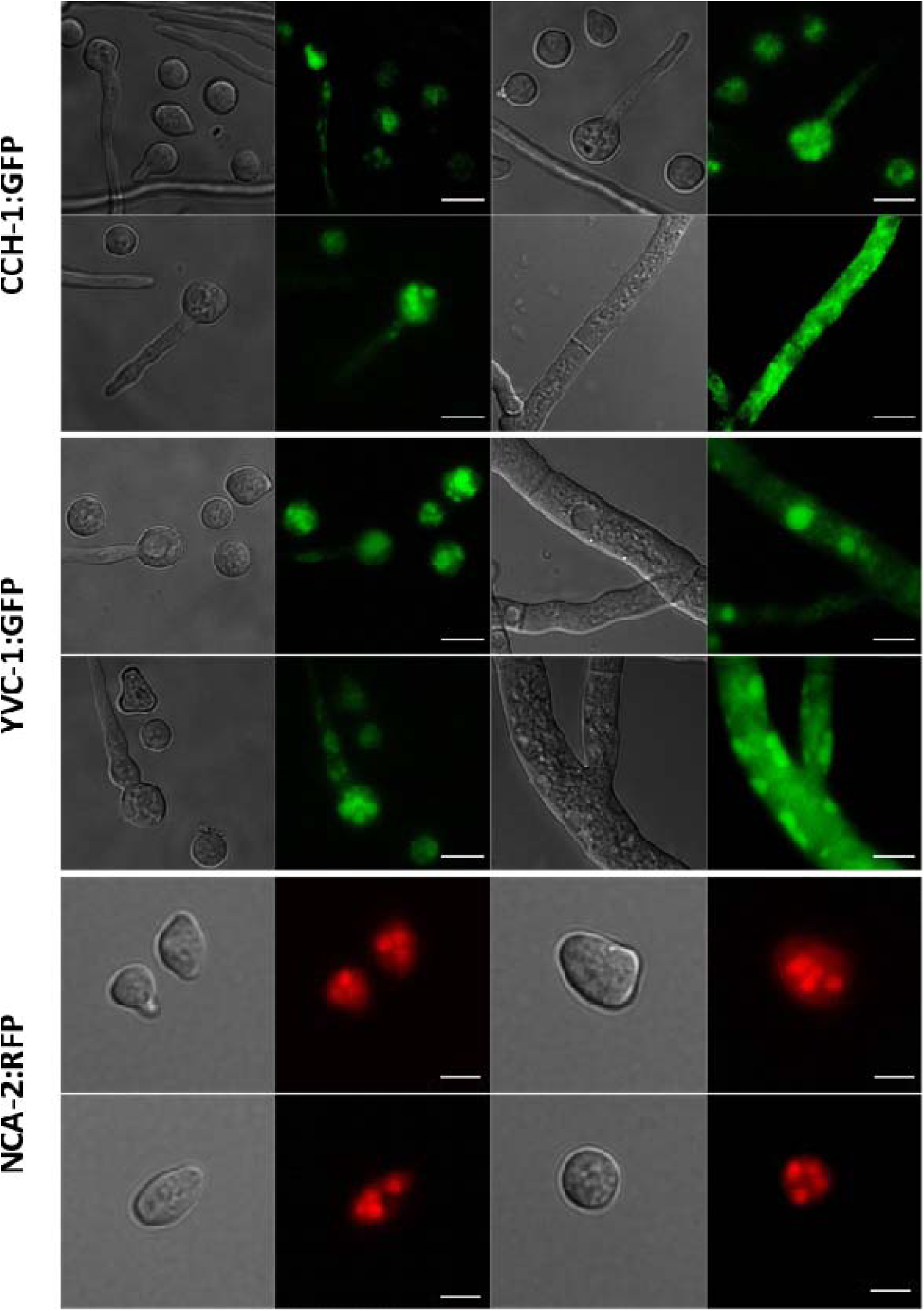
Localization of CCH-1 and YVC-1 labelled with GFP in macroconidia, germlings and vegetative hyphae. Representative widefield fluorescence images of the different cell types of GFP tagged CCH-1 and YVC-1 are shown. As NCA-2 has been previously localised in hyphae, only macroconida re shown for this RFP expressing strain(Bowman, Draskovic *et al*. 2009). CCH-1 appears to be localised in the vacuolar system in macroconidia and germlings, but also appears as distinct points in the plasma membrane of mature hyphae. YVC-1 appears to localise to the vacuolar system in all cell types. RFP tagged NCA-2 localises to the vacuole in macroconidia. Bar = 5 μm.

In order to localise the CCH-1 and YVC-1 proteins, plasmids were constructed based on FJ457002 for N-terminal tagging and FJ457006 for C-terminal tagging (Honda and Selker 2009).

Both the C- and N-terminal YVC-1:GFP fusion proteins fluoresced, although the N-tagged protein showed a stronger more stable fluorescence. Fluorescence mainly appeared within vacuoles in conidia, germ tubes and mature hyphae (Fig. 6). The CCH-1:GFP fusion products produced very little fluorescence, both from N and C tagged versions. This appeared to be localised to vacuoles within conidia and intracellular membranes within older hyphae. Single punctate spots of GFP fluorescence could however be seen at the plasma membrane of mature hyphae, Fig.6.

## DISCUSSION

### Ca^2+^ plays important roles in the mode-of-action of PAF26

In this research, evidence for Ca^2+^ signaling and homeostasis having a significant role in the PAF26 mode-of-action was obtained initially by testing the PAF26 sensitivity of conidia grown in the presence or absence of calcium. When the level of calcium in the media is kept at a minimal level, tolerance increases. Conversely when the level is raised, tolerance decreases. This was confirmed with deletion mutants defective in different components of their cellular transport machinery of Ca^2+^. Deletion of most of the Ca^2+^ channels significantly increased the concentrations at which PAF26 inhibits fungal growth. The elimination of the high affinity Ca^2+^ plasma membrane channel CCH-1 and the vacuolar channel YVC-1 as well as the Ca^2+^ ATPase resulted in inhibitory concentrations of the peptide approaching the point at which passive membrane permeabilisation occurs. Ca^2+^ has been shown to be an important factor in the mechanistics of PMAP-23 against *Candida albicans*, the disruption results in reactive oxygen species (ROS) accumulation which triggers apoptosis (Kim and Lee 2019). This effect is also seen in the peptide CGA-N9 against *C. tropicalis* (Li, Chen *et al*. 2019). High levels of cytosolic free Ca^2+^ have also been shown to confer protection to *C. albicans* against the antimicrobial MUC7 12mer, thought to be by a resulting change in the cell membrane properties preventing peptide entry (Lis, Liu *et al*. 2010). It appears in this case that the elevated levels of cytosolic free Ca^2+^ are a stress response designed to maximise survival by initiating the endocytosis of PAF26. There are significant changes to the pattern of peptide cell interaction in the mutants and to the quantity of peptide the mutants take up. The mutations in the Ca^2+^ distribution machinery appear to disrupt the internalisation and accumulation of the peptide which in turn increases tolerance to PAF26. Deletion of *nca-2* appears to influence the binding of PAF26 to the cell envelope, the *Δnca-2* mutant has been shown to accumulate up to 10 fold more Ca^2+^ than wild type cells, suggesting that NCA-2 serves to remove Ca^2+^ from the cell (Bowman, Abreu *et al*. 2011). We found no significant difference in the resting level of [Ca^2+^[_cyt_ from the wild type however under unstimulated conditions. Interestingly however, the *Δnca-2* mutant has a membrane potential reversed from the wild type due to a lack of cell surface H^+^ATPase function (Hamam and Lew 2012). In *S. cerevisiae* the H^+^ATPase is one of the most abundant cell surface proteins (Bagnat, Chang *et al*. 2001) and makes up a significant amount (up to 10%) of the cell surface in *N. crassa* (Bowman, Blasco *et al*. 1981). PAF26 has been shown to cause a rapid depolarization of the membrane in wild type cells in an energy independent manner (Muñoz, Marcos *et al*. 2012). Given that PAF26 has a net positive charge, the membrane potential reversed *Δnca-2* should technically not be inhibited in PAF26 membrane binding from an electrostatic view. It is therefore possible to propose that PAF26 directly inhibits H^+^-ATPase action, possibly by direct binding; misfunctioning H^+^--ATPase at the plasma membrane is sent to the vacuole for degradation in yeast, a possible mechanistic for the accumulation of PAF26 (Bagnat, Chang *et al*. 2001, Liu, Sitaraman *et al*. 2006). When the peptide is internalised in the *Δyvc-1* mutant, it remains in small vesicles with very little sign of the peptide entering the vacuolar system; adding support for the hypothesis that YVC-1 and a threshold amount of PAF26 is required to initiate vacuolar fusion. Vacuolar fusion occurs through conformational change of the docking SNARE proteins, triggered by the release of Ca^2+^ from the vacuole in *S. cerevisiae* (Bayer, Reese *et al*. 2003, Merz and Wickner 2004, Coonrod, Graham *et al*. 2013). In *S. cerevisiae*, isolated vacuoles are able to catalyse their own fusion through the release of Ca^2+^ from yvc1p present in the vacuolar membrane (Peters and Mayer 1998). This does not appear to be the case in *N. crassa* however, as both deletion of YVC-1 and removal of extracellular Ca^2+^ resulted in the trapping of the peptide in small vesicles. Therefore extracellular Ca^2+^ is required to initiate the release of Ca^2+^ from the vacuoles and trigger fusion. This raises questions as to why internal membrane fusion is reliant on external stimuli. In the *Δcch-1* mutant the peptide was directly translocated across the plasma membrane into the cytoplasm in an energy dependent manner, before accumulation in vacuoles, meaning PAF26 does not kill by being present in the cytosolic space. CCH-1 is therefore required to initiate the endocytic pathway. Energy dependent peptide uptake into the vacuolar system is also seen in the *Penicillium chrysogenum Penicillium* antifungal protein B (PAFB), where the fungal cells show no signs of cell death as long as the peptide remains in the vacuole (Huber, Hajdu *et al*. 2018). This is similarly seen in the *Neosartorya (Aspergillus) fischeri* antifungal protein (NFAP), which is also accumulated in the vacuole in an energy dependent maner. Again, this peptide also does not exert its antifungal effects until it is present within the cytoplasm (Hajdu, Huber *et al*. 2019). This contrasts the findings here of PAF26 being present within the cytoplasmic space without killing the cells. The importance of the vacuole in the mode of action of many antifungal peptides is becoming increasingly apparent. A genetic screen of *S. cerevisiae* revealed tolerance to the plant antifungal defensins NbD6 and SBI6 could be increased by deletion of genes involved in vacuolar transport (Parisi, Doyle *et al*. 2019).

Ca^2+^ measurement at the population level revealed that the Ca^2+^ responses to PAF26 were markedly different in the PAF26 tolerant mutants. Whilst all underwent a similar initial increase in [Ca^2+^]_cyt_ the second biphasic increase seen in the wildtype was abolished. This initial increase was independent of CCH-1, as the signal in the Δ*cch-1* mutant increased, but was dependent on external Ca^2+^ and L-type Ca^2+^ channels. The fact that L-type channel blockers completely stop the [Ca^2+^]_cyt_ increase as does the chelation of external Ca^2+^ using BAPTA would suggest that there is another, as yet unknown cell surface Ca^2+^ channel. These findings mirror those of Binder *et al*., 2010, who found that deletion of *cch-1* did not prevent the increase in [Ca^2+^] _cyt_ in response to *P. chrysogenum Penicillium* antifungal protein (PAF). When these measurements were taken at the level of the individual, rather than the population, it became apparent that PAF26 does not in fact cause a dose dependent rise in [Ca^2+^]_cyt_ but rather increases both the amplitude and frequency of Ca^2+^ spiking, again dependent on the presence of external Ca^2+^. These findings have allowed us to propose the model shown in Fig. 7 for the role of Ca^2+^ signalling and homeostasis in the mode of action of PAF26. Now the challenge is to identify the trigger of vacuolar release of PAF26 and the initiator of cell death.

**Fig 7.**
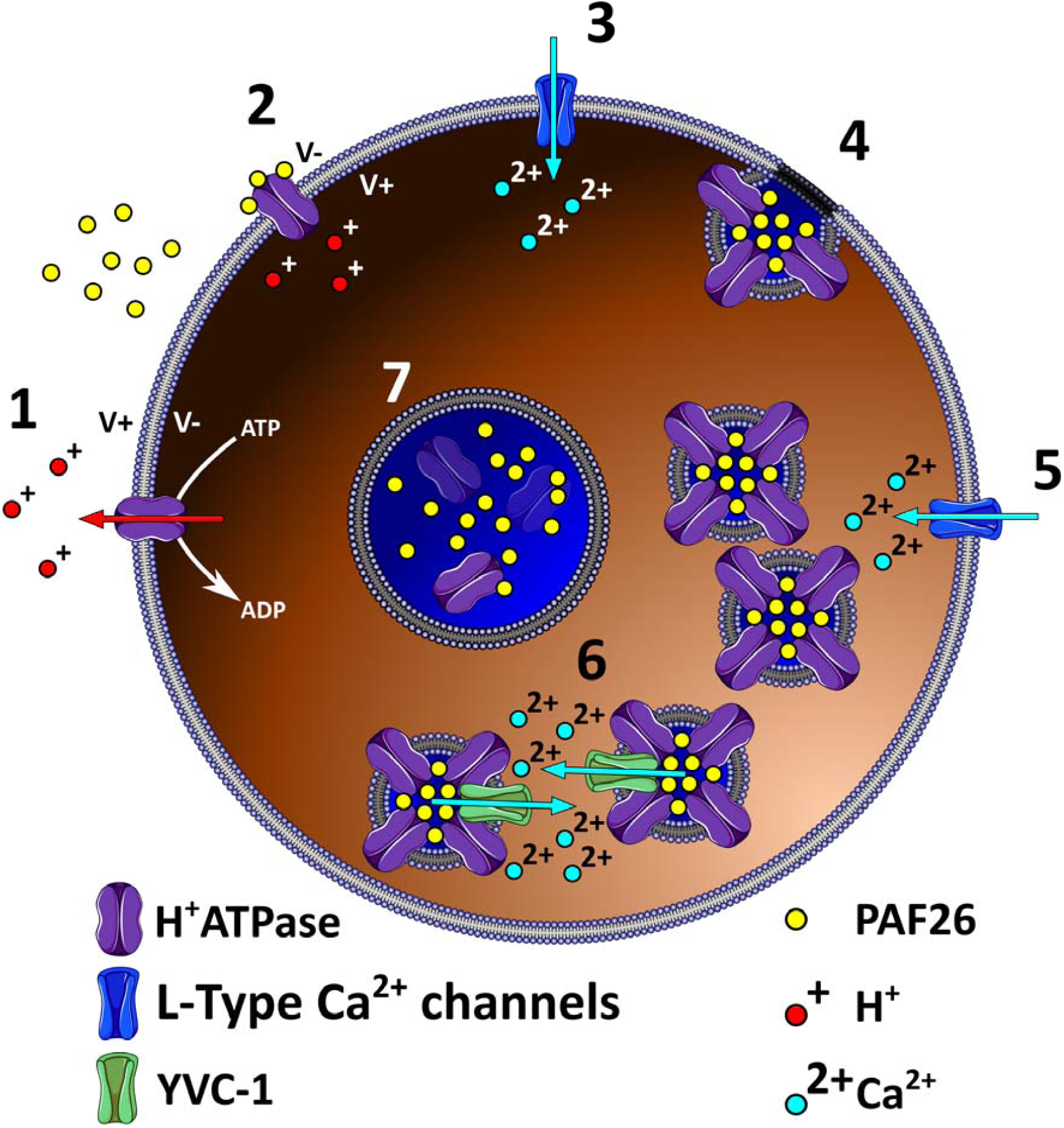
Proposed role of Ca^2+^ signalling in the mode of action of PAF26. **1** initially the cell is unstimulated and maintains a low [Ca^2+^]_cyt_ and stable membrane potential. **2** PAF26 inhibits the cell surface H^+^ATPase initiating membrane depolarisation. **3** Depolarisation of the plasma membrane activates the voltage gated CCH-1, causing [Ca^2+^]_cyt_ to rise. **4** This rise in [Ca^2+^]_cyt_ initiates the turnover of the H^+^ATPase including bound PAF26 by endocytosis. **5** further rises in [Ca^2+^]_cyt_ caused by external Ca^2+^ entering through CCH-1 initiate Ca^2+^ release through YVC-1 **6** driving the fusion of membranes. **7** PAF26 is released from the H^+^ATPase as it is degraded and accumulates in the vacuolar system.

## Materials and methods

### *Neurospora crassa* stocks

*N. crassa* strains were obtained from the Fungal Genetics Stock Centre (FGSC, University of Missouri, Kansas City, USA), unless otherwise specified. FGSC2489 (*matA*,74-OR23-1VA) was used as the wild type control. *N. crassa* strains generated in this study were derived from FGSC9717 (*matA*,Δ*mus-51::bar*^*+*^; *his-3*^*-*^), FGSC9720 (*matA*,Δ*mus-52::bar*^*+*^; *his-3*^*-*^) and FGSC6103 (*matA,his-3*^*-*^) *N. crassa* was cultured and maintained according to FGSC protocols (www.fgsc.net/Neurospora/NeurosporaProtocolGuide.htm) unless otherwise stated.

For microscopy, conidia were cultured in liquid VM for up to 4 hours at 30°C using 8 well Nunc™ Lab-Tek™ II Chambered Coverglass.

### *N. crassa* transformation

Electroporation of macroconidia was carried out using the following settings: resistance 600 Ω, voltage 1.5 kV/cm and capacitance 25 μFd.

### Heterokaryon purification

*N. crassa* strains that were obtained as heterokaryons, were purified to homokaryons by standard procedure (www.fgsc.net/Neurospora/NeurosporaProtocolGuide.htm).

### Plasmids

In order to create pTef1AeqS, aequorin was amplified from pAB19 using the primers AeqS-F and AeqS-R. The vector pCC019 was digested with PacI and EcoRI and the vector gel purified. Aequorin was then amplified with AeqS-IF-Fw and AeqS-IF-Rv to add overlaps homologous to the backbone whilst maintaining digestion sites and the vector was assembled using Gibson assembly. To generate pYVC1CGFP and pYVC1NGFP, the *yvc-1* ORF was amplified from gDNA using the primers NC-YVC1-AMP-FW and NC-YVC-1-AMP-RV. The purified product was then amplified using NC-YVC1-N-FW & NC-YVC1-N-RV for N terminal tagging and NC-YVC1-C-FW & NC-YVC1-C-RV for C terminal tagging. The FJ457002 vector was digested using PacI and XbaI and the FJ457006 vector digested with AscI and NotI. Following gel extraction both vectors were assembled using Gibson assembly.

Tagging of CCH-1 was not straightforward due to the size of the ORF - 6.5kb. The CCH-1 ORF was amplified first using the primer sets NC-CCH1 AMP-FW and NC-CCH1-3.5Rv and NC-CCH1-3.5Fw and NC-CCH1-AMP-RV to amplify the gene in two segments, 5’ and 3’. The complete constructs were built using fusion PCR; primer NC-CCH1-C-FW, 5’ segment, 3’ segment and primer NC-CCH1-C-RV, the N terminal tagging vector used the primers NC-CCH1-N-FW and NC-CCH1-N-RV. Following successful fusion the vectors were assembled as for YVC-1. Primer sequences are given in supplementary material.

### IC_50_ values

IC_50_ values were calculated using clear, U bottom polystyrene microtitre plates. PAF 26 was prepared at two concentrations, 35 µM and 30 µM, and diluted two-fold appropriately in to reach the final experimental concentrations. Conidia were diluted to a concentration of 1 × 10^6^ in 20% liquid VM and 50 µl dispensed into each well. Experimental PAF26 concentrations (µM) were: [17.50][15.00][8.75][7.50][4.38][3.75][2.19][1.88][1.09][0.94][0.55][0]. Final conidial concentration was of 5 × 10^5^ in 10% VM. Absorbance measurements were obtained at 610 nm. This wavelength is close to the 595nm at which the relationship between optical density and fungal biomass is linear (Broekaert, Terras *et al*. 1990).

### Calcium measurements

Coelenterazine (#C-7001, www.biosynth.com)was prepared by dissolving in ice cold methanol (MetOH) under a protective atmosphere in complete darkness to a concentration of 10 µg/µl. Stocks were diluted to the working concentration of 3.175 µg in 10 µl MetOH, 74.78 µM.

Conidia were diluted to a concentration of 1 × 10^6^ in 10% VM, coelenterazine was added to a final concentration of 2.5 µM. 100 µl of conidial suspension was used per well of a white microtitre plate and the plates incubated at 25°C in the dark for 6 hours. The light output of aequorin was measured by counting the photons emitted by a single well over one second, each well in a row of six being measured once every cycle of 7 seconds. After recording baseline luminescence, a single row was treated with PAF26 and the light output measured over time. Following this the second row was run with 100 µl 3M CaCl_2_:20% ethanol (EtOH) injected on cycle 8 to immediately discharge all the aequorin.

### [Ca^2+^] calculation

The raw relative light units (RLU) were converted to [Ca^2+^] using the formula:(Bonora, Giorgi *et al*. 2013)

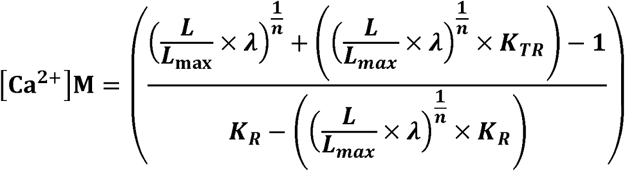

Where:

- ***L*** is the light intensity at sampling time
- ***L***_***max***_ is the total light emitted at sampling time
- **K**_**R**_ is the constant for calcium bound state **(7**,**230**,**000)**
- **K**_**TR**_ is the constant for calcium unbound state
- **(120)**
- **λ** is the rate constant for aequorin consumption at saturating [Ca^2+^] **(1)**
- **n** is the number of Ca^2+^ binding sites **(2**.**99)**

In order to calculate L_max_ the total RLU (RLU_total_) available was calculated from the discharge of aequorin (RLUd) using the trapezoid rule formula, where the total RLUd available at time point t_n_ equals:

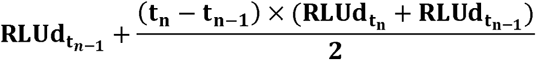

and RLUd_total_ is the final value reached. Switching to the experimental data (RLUe), L_max_ was calculated for each time point as:

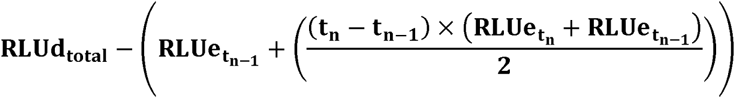

### Peptides and peptide handling

Peptides were ordered from GenScript (www.genscript.com) at > 98 % purity. Stock solutions were prepared in 50:50 dimethyl sulfoxide (DMSO):H_2_O buffer. Working peptide solutions were prepared in ddH_2_O.

### Microscopy

All microscopy was carried out on a Nikon Eclipse^®^ TE2000-E inverted microscope, using a Nikon Plan Apo 60 × 1.2 N.A. DIC H water immersion objective and Nikon G-2A and B-2A filter sets, excitation was provided by a CoolLED illumination system set at 550 nm for use with the G-2A filter or 470 nm for use with the B2-A filter.

Confocal microscopy was carried out using a Leica TCS SP8 equipped with two hybrid GaAsP detectors (HyD) and two photomultiplier tubes (PMT). Excitation was provided using either the Leica tuneable white light laser (450 −750 nm), an argon laser (458nm, 476nm, 488nm and 496nm) or UV laser (405nm). Images were captured using LAS X software and the Leica 63 X water immersion objective. All image handling and analysis was carried out using Fiji (fiji.sc/Fiji).

### Quantification of Ca^2+^ signalling

Fluorescence values were exported from Fiji and analysed in Excel. Data was normalised using feature scaling. This scales the data to removes irregularities in GCaMP6 expression and photon yield. The function used was: *x*^I^ = MIN + (*x*-*min*_*x*_)(MAX-MIN)/(*max*_*x*_-*min*_*x*_). Where MIN and MAX are the required scale range, *x* is the original value and *x*^*I*^ is the normalised value. *min*_*x*_ is the minimum value for that data set and *max*_*x*_ is the maximum, *max*_*x*_ was defined as the maximum value from all experiments plus 10 to avoid artificial amplification of noise. The noise was removed using an IF function: =IF(*x*>*y,x*,0) where *x* is the data point and *y* is the noise threshold limit. Finally, an IF(AND(*x*>*x*_-*1*_, *x*>*x*_*+1*_),1,0) function, where *x*_*-1*_ and *x*_*+1*_ are the surrounding measurements, quantified each peak in GCamP6 fluorescence

### Data availability

Data used in this study is available at: http://dx.doi.org/10.17632/sb2r2zsxdz.1

## Supplementary

### Primers

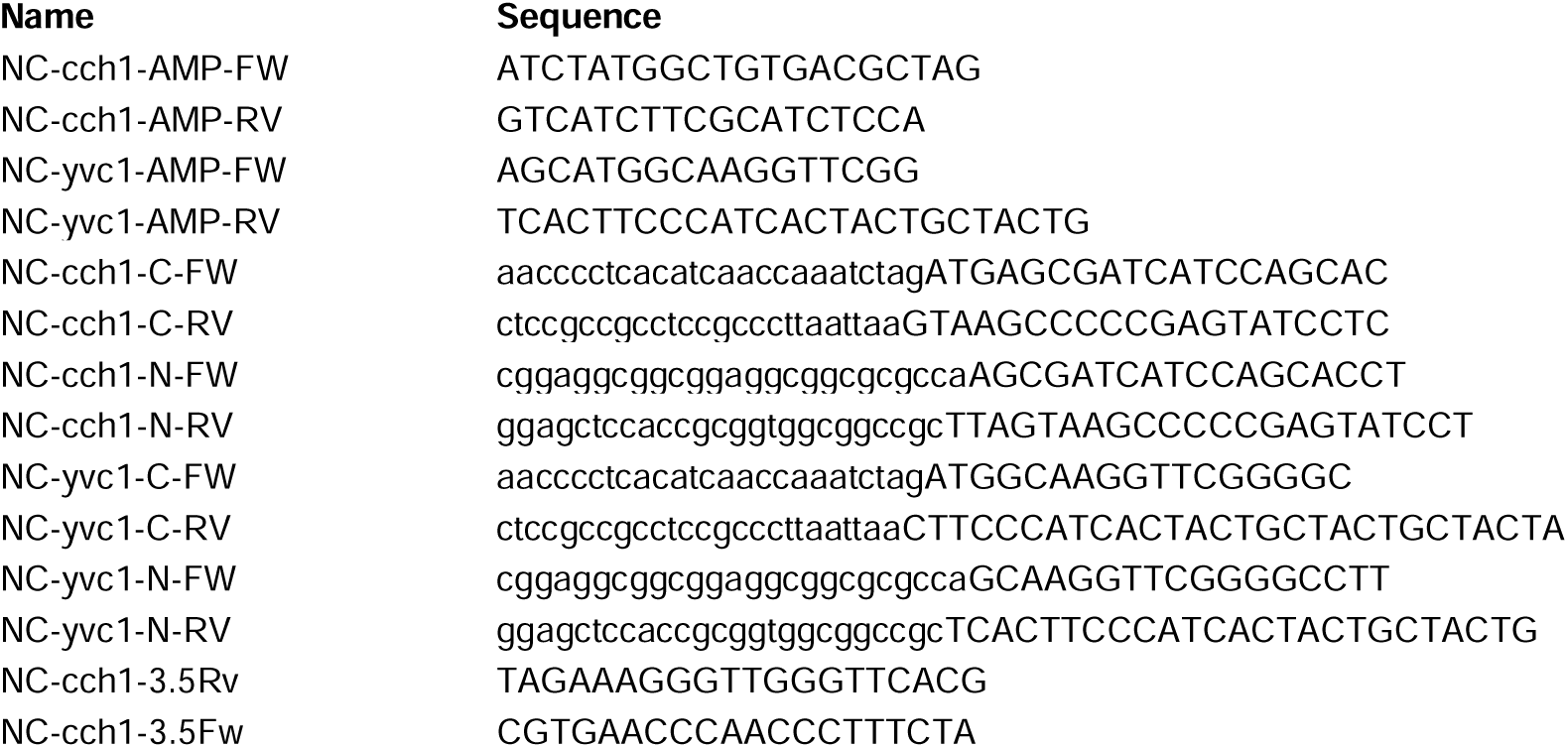

### Plasmids

| Plasmid | Backbone | Genotype | Source |
| --- | --- | --- | --- |
| FJ457002 |  | *Pccg-1::c-gly::gfp* | Honda & Selker, 2009 |
| FJ457006 |  | *Pccg-1::n-gfp* | Honda & Selker, 2009 |
| pAB19 | pAZ6 | *Pccg-1-AeqS-PtrpC-bar<sup>+</sup>* | Marris, 2007 |
| pCC019 | FJ457002 | *his-3-Ptef-1-gcamp6s* | Chang, 2015 |
| pTef1AeqS | pCC019 | *his-3-Ptef-1-AeqS* | This study |
| pYVC1CGFP | FJ457002 | *his-3-Pccg-1-yvc-1-gfp* | This study |
| pYVC1NGFP | FJ457006 | *his-3-Pccg-1-gfp-yvc-1* | This study |
| pCCH1CGFP | FJ457002 | *his-3-Pccg-1-yvc-1-gfp* | This study |
| pCCH1NGFP | FJ457006 | *his-3-Pccg-1-gfp-yvc-1* | This study |

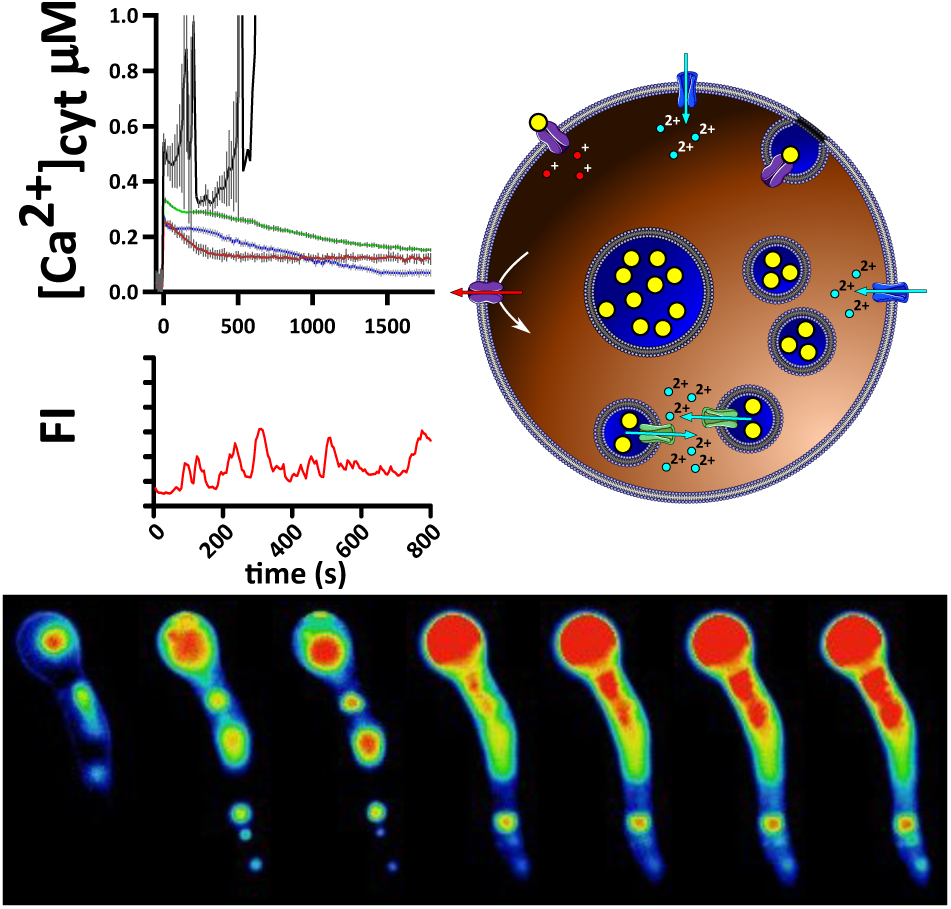

